# Profibrotic Changes Following Tension Application in a Fetal Lamb Model of Long Gap Esophageal Atresia

**DOI:** 10.64898/2026.01.26.701811

**Authors:** Jessica C. Pollack, Nicolas Vinit, Shelley Jain, Rachel Conan, Melanie Bates, Mia Kwechin, Alicia Eubanks, Mike Xie, Amanda Muir, Emily Partridge

## Abstract

**Introduction:** Esophageal atresia is a common congenital anomaly, occurring in 1 in 3,500 live births. The Foker process has revolutionized the treatment of long gap esophageal atresia (LGEA). It is well established that the Foker process causes tension accelerated growth of the esophagus, but what occurs at the molecular level during tension accelerated growth is still unknown. We aimed to create tension accelerated growth in a fetal lamb model of LGEA in order to answer this question.

**Methods:** Following IACUC approval, time-dated fetal lambs (108 to 120 days of gestation) underwent thoracic esophagectomy. Both esophageal ends were ligated and sutured together to create an internal pexy under high tension. Lambs were delivered on postoperative day 2 (POD2) (n=7), POD6 (n=9) or term (n=5). The native esophagus collected at model creation served as control tissue. Specimens were bluntly separated into two layers: inner layer (IL) (epithelium, lamina propria, muscularis mucosa, submucosa) and outer layer (OL) (submucosa, muscle layer, adventitia). RNA sequencing (RNAseq), proteomics, immunohistochemistry, western blotting and real-time qRT-PCR were performed on the specimens. Mann-Whitney’s or unpaired t-test were used for statistical analyses. Esophageal fibroblast cell lines established from human biopsy specimens were cultured and stimulated with TGF-beta for in vitro studies on collagen expression.

**Results:** 23 lambs underwent esophagectomy with tension suture placement at 108 to 120 days gestation. Histologic analysis of tension conditioned compared to control esophagus by trichrome staining demonstrated an increase in collagen deposition in tension conditioned esophagus compared to controls. High throughput bulk RNA sequencing and proteomic analysis were performed with a focus on pathways implicated in fibrosis. GSEA analysis of the inner layer demonstrates upregulation of TGFB signaling, extracellular matrix organization, and collagen deposition at all timepoints. Further analysis was performed to evaluate specific collagen subtypes contributing to this profibrotic phenotype, and COL8A1 and COL12A1 were both significantly upregulated in both RNA and proteomic analysis at all timepoints, with Western blotting confirming up regulation in stretched tissue. In order to evaluate the relationship between TGFB signaling and collagen deposition in the esophagus, we stimulated esophageal fibroblasts with TGFB, qRT-PCR was performed to evaluate the expression of COL8A1, COL12A1, and COL6A3. Expression of all three of these collagen subtypes was noted to be significantly upregulated at all timepoints following TGFB stimulation when compared to non-stimulated controls.

**Conclusions:** Tension accelerated growth can safely be achieved in a fetal ovine model of long gap esophageal atresia. Additionally, esophageal atresia can be modeled in the ovine fetus as early as 92 days gestation. Our results demonstrate that esophageal tissue subjected to sustained tension undergoes significant profibrotic changes, as evidenced by upregulation of TGFB signaling, alterations in extracellular matrix organization, and increased collagen deposition. While it is well documented that patients with LGEA have an increased risk of post operative esophageal strictures, these findings provide the first in vivo proof of the role of tension in conferring a profibrotic phenotype in the tension-lengthened esophagus.

## Introduction

Esophageal atresia (EA) is a common congenital anomaly that results in discontinuity of the esophagus. EA is present in 1 in 3500 live births (1) and the standard treatment consists of a primary anastomosis, or surgical connection, of the proximal esophagus to the distal esophagus. In some cases, the gap between the proximal pouch and distal pouch is too large, and primary repair is not possible. These cases are referred to as long gap esophageal atresia (LGEA) (2). The current treatment for LGEA is based on the concept of tension accelerated natural growth. To achieve this, the blind esophageal ends are placed on tension that is gradually increased over time to cause longitudinal tissue growth (3). This growth decreases the gap between the proximal and distal esophageal pouches, thus allowing for delayed primary repair (4). This process is referred to as the Foker procedure and it is typically performed over the first weeks to months of life.

LGEA is associated with significant morbidities including prolonged hospital length of stay, increased total ventilator days, increased risk of anastomotic leak, and high rates of post-operative stricture formation (5–7) compared to patients in whom primary repair is achieved. While the Foker procedure has significantly advanced the management of LGEA, it has not eliminated the morbidity experienced by this patient population. One of the most common complications following EA repair, particularly in cases of LGEA, is esophageal stricture. This is defined as a narrowing of the esophagus that often occurs at the site of anastomosis (8,9). Strictures cause difficulty with swallowing and lead to significant long-term morbidity, often requiring alternative feeding access and multiple procedures under anesthesia to dilate the narrowed segment of esophagus. Little is known about the molecular mediators involved in stricture formation, there are no known methods for prevention, and there are no non-invasive techniques for treatment.

A robust animal model is imperative to improve our understanding of tension accelerated natural growth, improve upon current surgical interventions and create a platform to study new surgical and therapeutic interventions that further minimize the morbidity associated with LGEA. Several studies have been performed using rodent models of EA, and while these models have generated valuable insight into the disease process, their small size limits the applicability to translational studies (10,11). Studies on translational methods in esophageal lengthening have utilized piglets given their larger size (12–14). However, this model is created at 8-12 weeks of age, while EA develops during embryologic development in humans. It has also been shown that while tension accelerated natural growth of the esophagus is possible in the neonatal period, it is ineffective in adults. Ideally, a robust model requires creation of the esophageal defect during gestation to closely mimic the human course of disease and allow for tension accelerated natural growth early in life.

The creation of a fetal animal model of LGEA has multiple benefits. It creates the opportunity to evaluate prenatal interventions for LGEA, as up to 40% of cases of EA are diagnosed prenatally (15), with rates as high as 85% in LGEA specifically (16). A fetal animal model would also eliminate the need to create an alternative method of nutrition during an extended period of esophageal discontinuity, given that the fetus would continue to be supported by the mother. The fetal lamb has long been regarded as the optimal animal model for fetal intervention given its similarities in size and anatomy to humans throughout gestation. In addition, the unique placentation, with multiple cotyledons as opposed to a single placenta, makes the ovine uterus highly tolerant to open fetal surgery at a wide range of gestational ages, and the fetal lamb models of congenital diaphragmatic hernia (17) and myelomeningocele (18) were both foundational in the development of prenatal and postnatal therapies.

In this manuscript, we describe the creation of a fetal lamb model of long gap esophageal atresia and the implementation of a tension procedure that aims to mimic the Foker procedure. We demonstrate that this procedure leads to profibrotic changes in the esophageal tissue.

## Methods

Animal experimental protocols were approved by the Children’s Hospital of Philadelphia Institutional Animal Care and Use Committee (Protocol #22-001467). Time-dated pregnant ewes from an approved vendor were used.

### Model creation

Time-dated pregnant ewes at 108 to 120 days gestation (term = 145 days) were anesthetized using inhaled isoflurane. The abdomen was entered through a midline laparotomy to expose the gravid uterus. Hysterotomy was performed and the right chest of the fetus was exposed. A posterolateral thoracotomy was performed, and the lung was retracted to visualize the esophagus. Dissection was performed to mobilize the esophagus from the level of the thoracic inlet to the diaphragm. The esophagus was ligated proximally and distally using 0 silk ties and the entire intrathoracic esophagus, between the two silk ties, was excised. 0 silk sutures were used to approximate the proximal and distal ends of the esophagus under maximal tension achievable without compromising tissue integrity. The thoracotomy was closed, and the fetus was returned to the uterus. The hysterotomy and laparotomy were closed.

### Tissue collection

Normal esophagus was collected at the time of model creation to serve as control tissue. Lambs were delivered by cesarean section on post operative day (POD) 2, POD 6, or near term (approximately 135 days gestation). Collection at multiple timepoints was performed to allow for tissue analysis throughout the process of tension accelerated growth. Collection near term was used to assess for survival following a fetal intervention for LGEA. After delivery, lambs were euthanized, and the esophagus was harvested. A full thickness section was fixed in formalin for histological analysis. The remainder of the esophagus was separated into an inner mucosal layer and outer muscular layer and representative sections were placed in RNA later solution for RNA sequencing analysis, all protect solution for western blot, and snap frozen for proteomic analysis.

### Histology

Sheep esophagus samples were fixed in 10% Neutral Buffered Formalin for 24 hours and submitted for standard paraffin processing. H&E staining was performed on a Gemini AS Automated Slide Stainer (Epredia). Masson’s trichrome staining was performed manually using standard protocols (19). Slides were cover slipped and scanned at 20x magnification with an Aperio AT2 Digital Slide Scanner (Leica Biosystems). Trichrome quantification was performed using Colour Deconvolution (20,21) software (version 2.1) in ImageJ (version 1.54).

### RNA sequencing

RNA was extracted from esophageal tissue samples using the RNeasy mini kit (Quiagen) according to manufacturer’s instructions. RNA-seq was performed using whole transcriptome 50 base pair single-end sequencing on Illumina HiSeq2500. STAR software suite was used to perform alignment, Subread’s feature counts for annotation, and R software packages including Seurat and Monocle for statistical analyses. Raw short read files were quantified using Kallisto (version 0.45.0) against the Ovis aries (OA) reference genome from NCBI. The output from Kallisto was then imported directly into DESeq2 (version 3.1) (22) to identify differentially expressed genes (DEGs). DEGs were defined as genes with a False Discovery Rate (FDR) of less than 0.05 and a fold change of 1.5. We annotated OA genes to their human homolog genes using NCBI Datasets command-line tools datasets and dataformat. All analyses were conducted using R (version 4.3). The volcano plot was created using the R package “EnhancedVolcano” (version 1.6). Gene Set Enrichment Analysis (GSEA) and plots were performed using GSEA (version 4.3.3) downloaded from the Broad Institute. To evaluate our RNA-Seq results using the GSEA tool, we applied ‘GSEApreranked’ tool as recommended to our ranked gene list and estimated gene set enrichments among the gene ranked list. All default parameters were applied.

### Proteomics

Snap frozen tissue samples of esophageal tissue were cryopulvarized followed by bead beating in SDS buffer for protein extraction. Samples were further processed for bottom up LCMSMS analysis using standard S-trap (Protifi) protocols (23) with Trypsin and LysC proteolysis. Tryptic peptides were analyzed by LCMSMS using Data Independent Acquisition (DIA) on a Thermo Exploris 480 or Bruker timsTOF pro2. The raw data was processed using Spectronaut (Biognosys AG) (24) against the uniprot Ovis aries reference proteome. System suitability of the mass spectrometers was monitored using QuiC software (Biognosys; Schlieren, Switzerland) for the analysis of spiked-in iRT peptides. As a measure for quality control, standard E. coli tryptic digest was injected in between samples (one injection after every five biological samples) and the data was collected in data dependent acquisition (DDA) mode. The collected DDA data was analyzed in MaxQuant (25) and the output subsequently visualized using the PTXQC (26) package to track the quality of the instrumentation.

An in-house generated R package was used for proteomics data processing and statistical analysis. The MS2 intensity values generated by Spectronaut were used to analyze the whole proteome data. The data was log2 transformed and normalized by subtracting the median for each sample. Data was filtered to have a complete value for a protein in at least one cohort. To compare proteomics data between groups, Limma t-test was employed to identify differentially expressed proteins, and volcano plots were generated to visualize the affected proteins while comparing different groups. Lists of differentially abundant proteins were sorted based on the adjusted P Value < 0.05, yielding a prioritized list for downstream bioinformatic analysis.

### Western Blotting

Tissue samples were homogenized in RIPA buffer and total protein concentration was quantified using the Pierce™ BCA Protein Assay Kit according to the manufacturer’s protocol. Protein samples were prepared with Pierce™ LDS Sample Buffer (Non-Reducing) and NuPAGE Sample Reducing Agent. Equal amounts of protein were loaded onto NuPAGE 3–8% Tris-Acetate gels and separated using NuPAGE Tris-Acetate SDS Running Buffer. HiMark™ Unstained High Molecular Weight Protein Standard was used as a molecular weight marker. Electrophoresis was performed until the ladder reached the bottom of the gel. Proteins were transferred onto nitrocellulose membranes using the Invitrogen™ iBlot™ 2 Gel Transfer Device. Membranes were blocked with 3% bovine serum albumin (BSA) in TBST (Tris-buffered saline with 0.1% Tween-20) for 90 minutes at room temperature. Membranes were incubated overnight at 4°C with primary antibodies against Col12a1 (1:50, Santa Cruz Biotechnology) and Col8a1 (1:1,000, Proteintech) prepared in 2% milk in TBST. Membranes were then incubated in GAPDH primary antibody (1:1,000, Cell Signaling Technology) for 90 minutes at room temperature. Following primary antibody incubation, membranes were washed and incubated for 45 minutes at room temperature with secondary antibodies: IRDye 800CW anti-mouse (1:20,000, LI-COR Biosciences) and IRDye 800CW anti-rabbit (1:20,000, LI-COR Biosciences). Protein bands were visualized using the LI-COR Odyssey 9120 Infrared Imaging System. Western blots were quantified using ImageJ (version 1.54).

### Fibroblast stimulation

Primary fetal esophageal fibroblasts (FEF3 cells) were seeded and cultured (Dulbecco’s Modified Eagle Medium (DMEM), 10% fetal bovine serum, 1% Penicillin, 1% Streptomycin) at 37°C for 24 hours, until reaching confluence, before stimulation with TGFB at 10 ng/mL (DMEM, 1% Penicillin, 1% Streptomycin) for 30 minutes, 1 hour, 2 hours, 4 hours, and 6 hours. Non-stimulated FEF3 cells were used as controls. The cells were then harvested to perform RNA extraction (RNeasy kit, Qiagen) and reverse transcription. Real-time quantitative RT-PCR was performed (TaqMan Gene Expression Assays) for COL8A1, COL12A1, COL6A3, and GAPDH. Relative mRNA levels of each gene were normalized to GAPDH levels for each condition and compared to controls using a paired t-test. A p-value <0.05 was considered significant.

## Results

### Creating a fetal lamb model of long gap esophageal atresia

In order to evaluate the impact of esophageal lengthening techniques, we created a fetal lamb model of long gap esophageal atresia. Complete intrathoracic esophagectomy was performed in fetal lambs during late gestation (108-120 d GA). The fetuses were then returned to the uterus to continue gestation. Delivery via cesarean section and necropsy were performed once animals reached near term (approx. 135 d GA). The procedure was well tolerated with no evidence of critical polyhydramnios, fetal loss, or preterm delivery.

One fetal lamb underwent intrathoracic esophagectomy during mid gestation (92 d GA) to evaluate if the procedure would be tolerated at an earlier gestational age. The procedure was well tolerated in this animal. The proximal esophageal pouch was noted to be significantly dilated when compared to the distal pouch.

### Performing an esophageal tension procedure in a fetal lamb model of long gap esophageal atresia

A series of pilot studies were performed to identify the optimal technique for placing tension on the proximal and distal esophageal pouches. Once pilot studies were complete, 19 fetal lambs underwent LGEA creation and tension procedure (Fig. 1) during late gestation (108-120 d GA). Total intrathoracic esophagectomy was performed, leaving two ligated blind ends of the esophagus. Next, tension sutures were used to reapproximate the proximal and distal esophageal pouches in order to achieve maximal tension without tissue damage. Fetuses were then returned to the uterus to continue gestation. Lambs were delivered by cesarean section and necropsy was performed at varying timepoints following the tension procedure in order to evaluate the effect of tension over the course of time. Seventeen fetal lambs were included in the study (Table 1) while two were noted to have intrauterine fetal demise (IUFD) at the time of delivery and tissue was not collected.

**Table 1.** Surgical details for all fetal lambs that underwent tension application during late gestation with survival to harvest.

| Animal Number | Experimental Group | Resection Length (cm) | Creation GA (days) | Necropsy GA (days) |
| --- | --- | --- | --- | --- |
| 6245 L1 | POD 2 | 4 | 113 | 115 |
| 6245 L2 | POD 2 | 3 | 113 | 115 |
| 6879 | POD 2 | 4 | 120 | 122 |
| 7062 | POD 2 | 3.5 | 120 | 122 |
| 5971 L1 | POD 2 | 3.5 | 120 | 122 |
| 5971 L2 | POD 2 | 3.5 | 120 | 122 |
| 5424 | POD 6 | 3.5 | 115 | 121 |
| 6726 | POD 6 | 3 | 108 | 114 |
| 6744 L1 | POD 6 | 3 | 108 | 114 |
| 6744 L2 | POD 6 | 3 | 108 | 114 |
| 2019 L1 | POD 6 | 3.5 | 120 | 126 |
| 2019 L2 | POD 6 | 3.5 | 120 | 126 |
| 7701 | Term | 3.5 | 113 | 135 |
| 7776 L1 | Term | 3 | 113 | 135 |
| 7776 L2 | Term | 3 | 113 | 135 |
| 6978 | Term | 4 | 120 | 135 |
| 6838 | Term | 2.5 | 120 | 133 |

**Figure 1.**
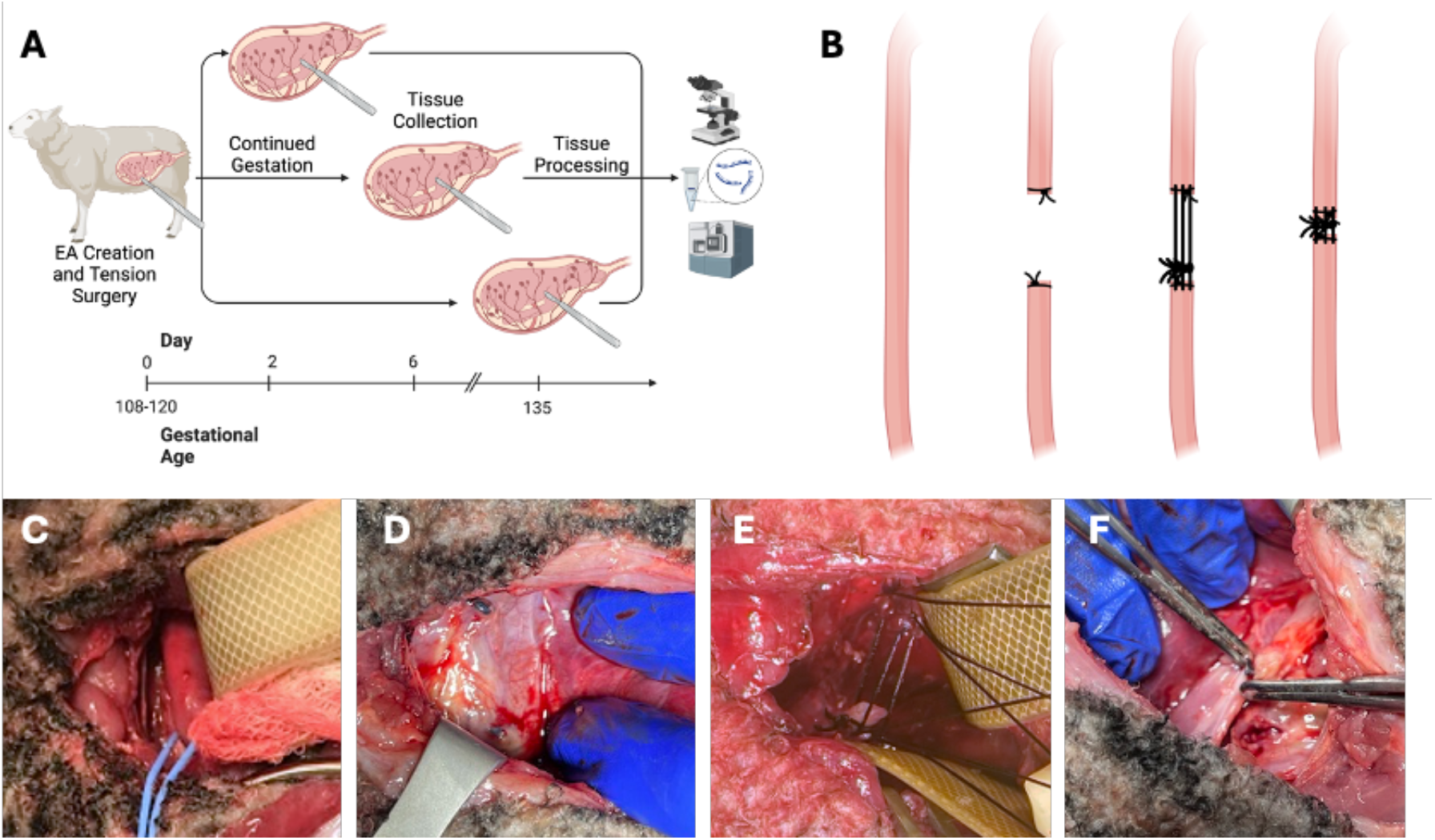
Creation of long gap esophageal atresia by intrathoracic esophagectomy in a fetal lamb. A: timing of surgery and tissue collection. B: schematic of surgical approach. C-F: Photographs of thoracic dissection with vessel loop surrounding esophagus (C), resection of entire intrathoracic esophagus with ligation of proximal and distal ends (D), tension sutures on proximal and distal pouches (E), and approximation of proximal and distal pouches with tension sutures (F).

Given the success of the LGEA model creation at an earlier gestational age, an attempt was made to perform LGEA creation and tension procedure during mid gestation (94 d GA) in two fetal lambs. The esophagus was noted to be extremely friable during the tension procedure and IUFD was noted at the time of delivery (134 d GA) for both lambs.

### Profibrotic changes present in tension conditioned esophagus

Histologic analysis of tension conditioned compared to control esophagus was performed to evaluate for pro-fibrotic changes occurring in the esophageal tissue throughout the course of tension application. H&E and trichrome staining was performed on tissue samples obtained at 2 and 6 days after tension application, as well as tissue collected near term (approx. 20 days after tension application). Trichrome staining demonstrated an increase in collagen deposition in tension conditioned esophagus compared to controls (Fig 2).

**Figure 2.**
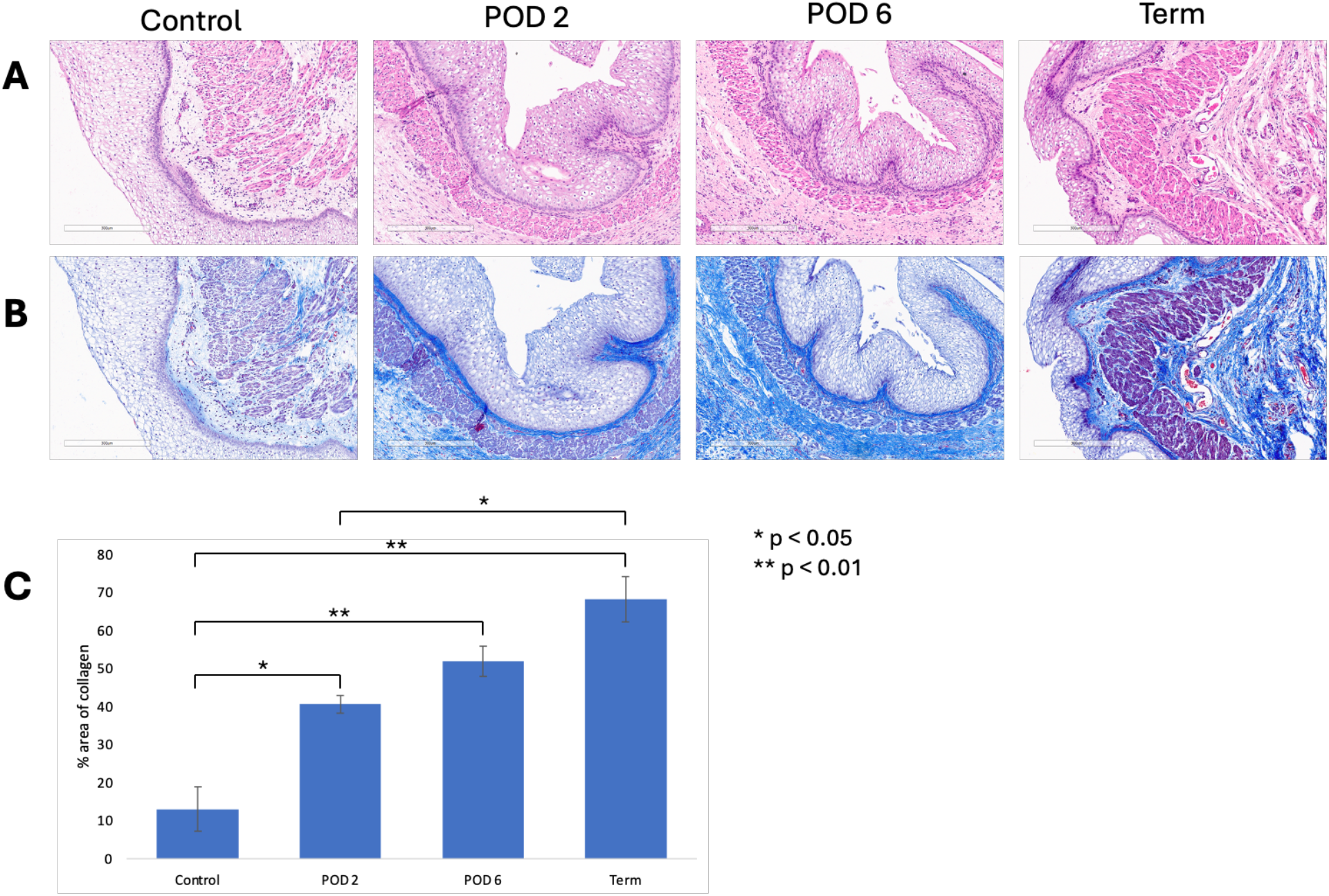
Histological analysis of esophageal tissue exposed to tension for 2, 6, and 14+ days compared to control. Visualized at 10x magnification. A: H&E stain. B: Masson’s Trichrome stain showing increased collagen deposition in the lamina propria of tension conditioned esophagus compared to control. C: Quantification of Masson’s Trichrome stain demonstrates increased collagen deposition in tension conditioned tissue compared to control.

Following RNA and protein extraction from the inner (mucosa/submucosa) and outer (muscularis) layer of esophageal tissue, high throughput bulk RNA sequencing and proteomic analysis were performed with a focus on pathways implicated in fibrosis. GSEA analysis of the inner layer shows robust upregulation of TGFB signaling, extracellular matrix organization, and collagen deposition at all timepoints (Fig. 3).

**Figure 3.**
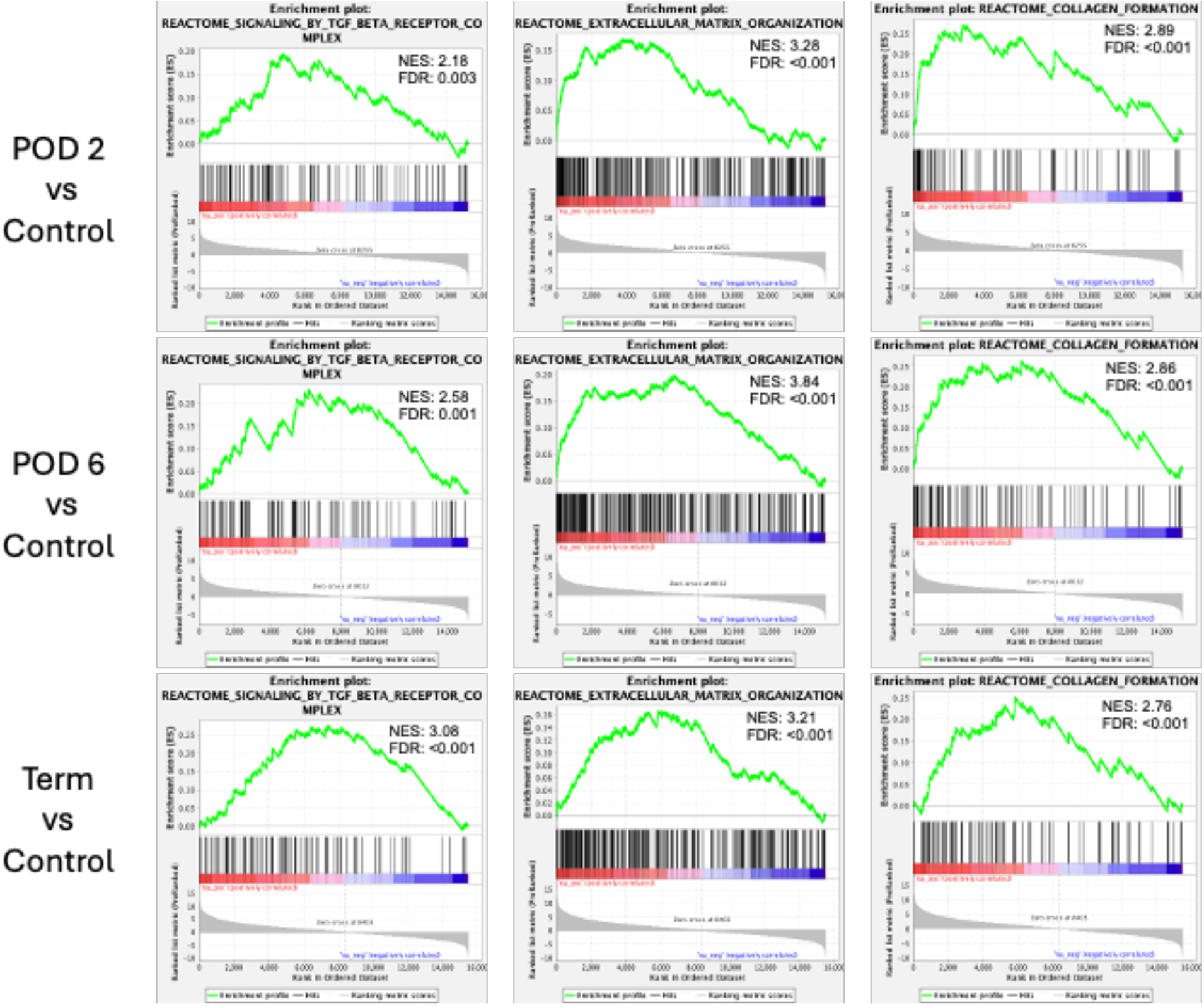
GSEA analysis of TGFB signaling, ECM organization, and collagen formation shows robust upregulation in tension-conditioned tissue across all timepoints compared to control tissue.

Given the upregulation in collagen formation, further analysis was performed to evaluate specific collagen subtypes. Fifteen collagen subtypes were significantly upregulated at one or more timepoints in tension conditioned esophagus compared to control (Fig. 4). Eight collagen subtypes had significantly higher protein concentration at one or more timepoints in tension conditioned tissue compared to control (Fig. 5). Of note, COL8A1 and COL12A1 were both significantly upregulated in both RNA and proteomic analysis at all three timepoints. Western blot also showed an increase in COL8A1 and COL12A1 in tension conditioned tissue compared to control (Fig. 6).

**Figure 4.**
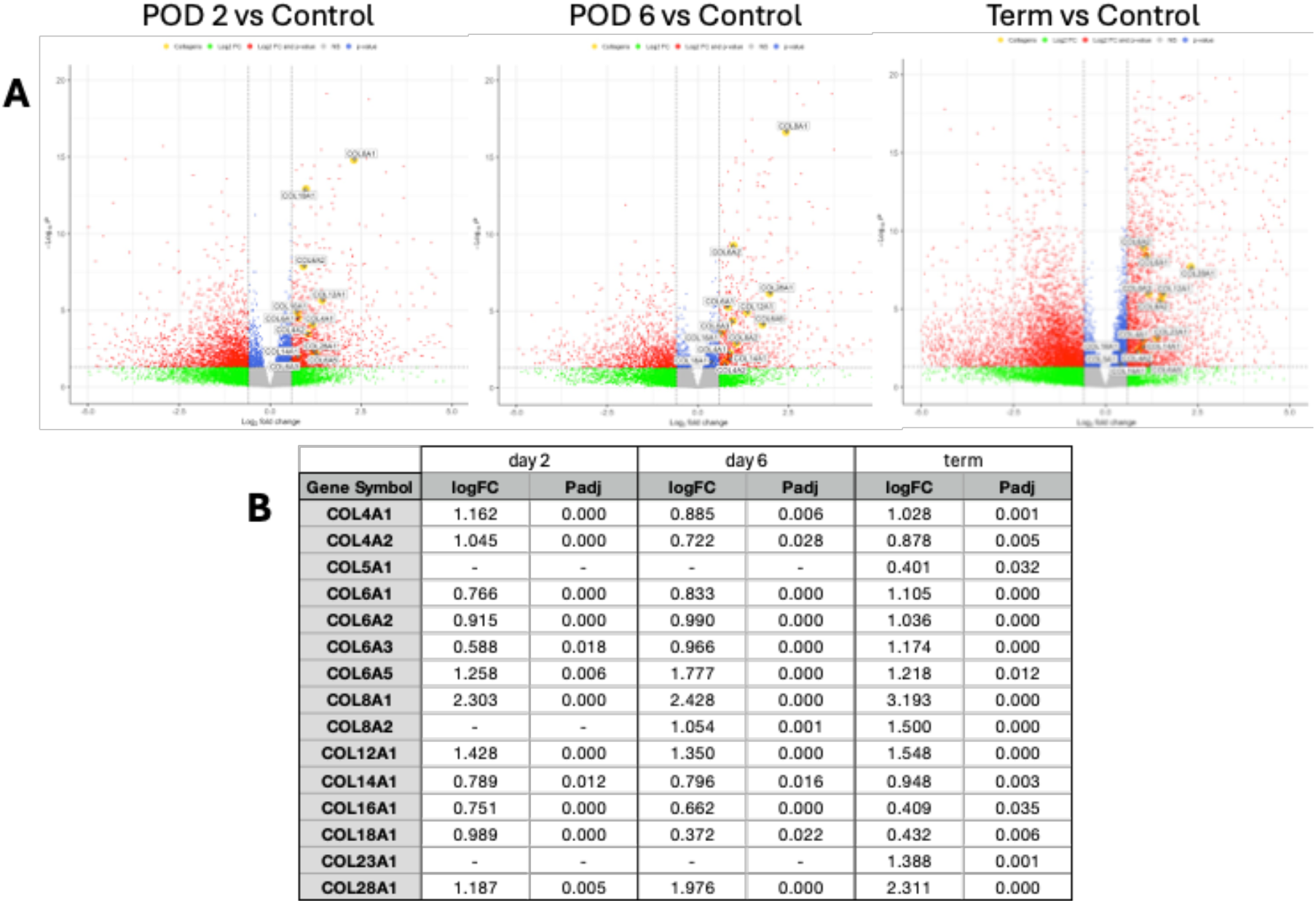
A: volcano plots showing upregulation of mRNA of specific collagen subtypes at 2, 6, and 14+ days following tension application when compared to controls. B: Tabulation of numerical values of relative mRNA expression.

**Figure 5.**
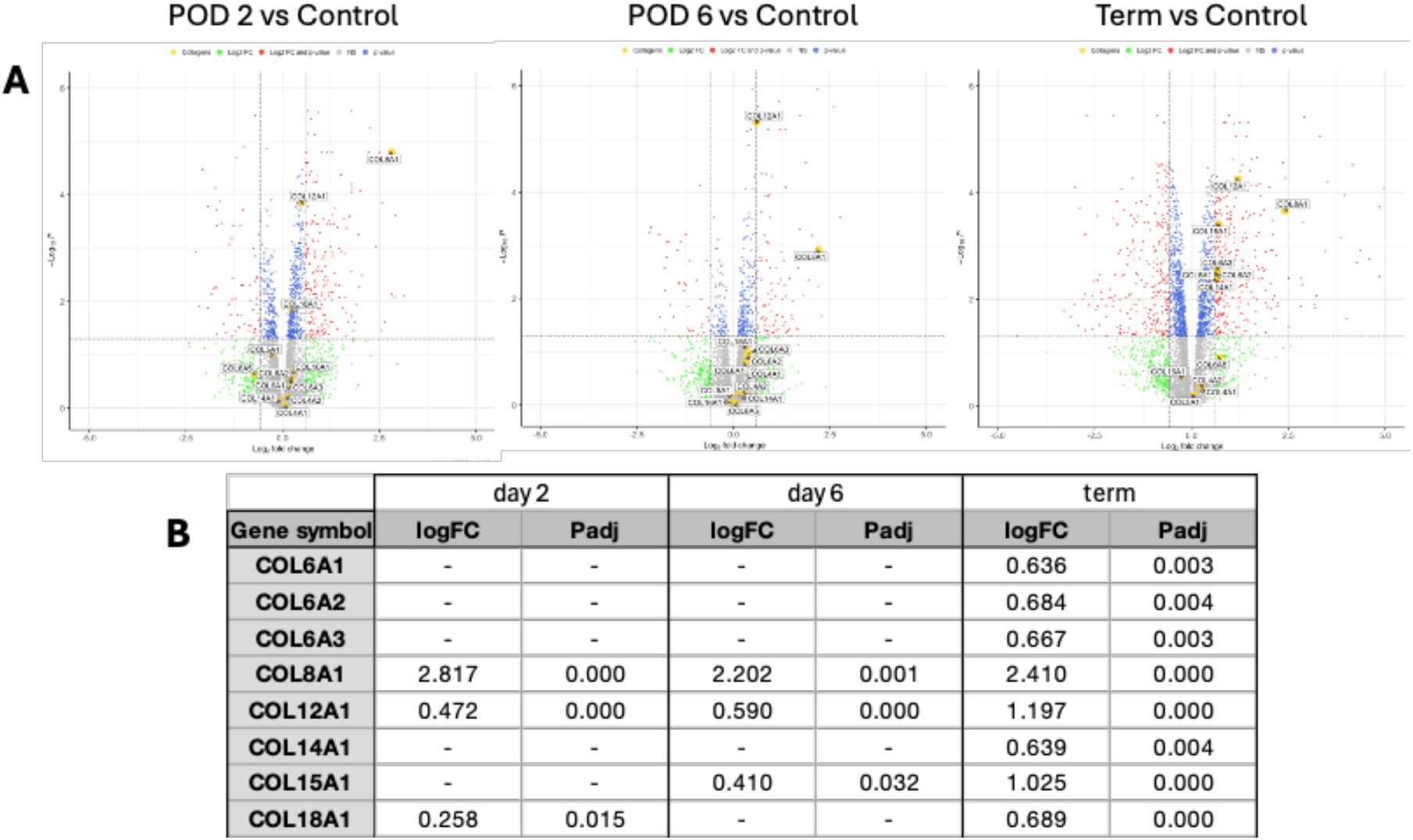
A: volcano plots showing increased protein concentration of specific collagen subtypes at 2, 6, and 14+ days following tension application when compared to controls. B: Tabulation of numerical values of relative protein concentration.

**Figure 6.**
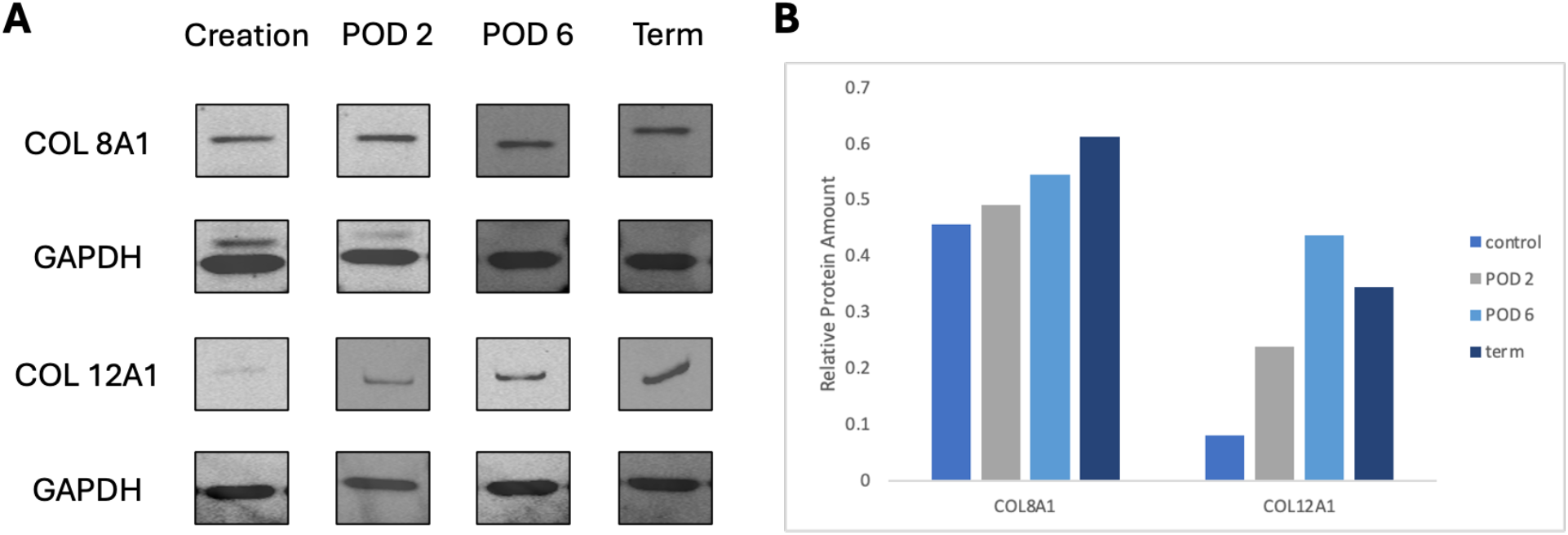
A: Representative western blots of COL8A1 and COL12A1 at each timepoint. B: Quantification of protein shows increased COL8A1 and COL12A1 in tension conditioned tissue compared to control.

In order to elucidate more specific potential targets in the profibrotic pathway, we evaluated the differential expression of interleukins with a focus on those that are upregulated in tension conditioned tissue compared to control (Fig. 7).

**Figure 7.**
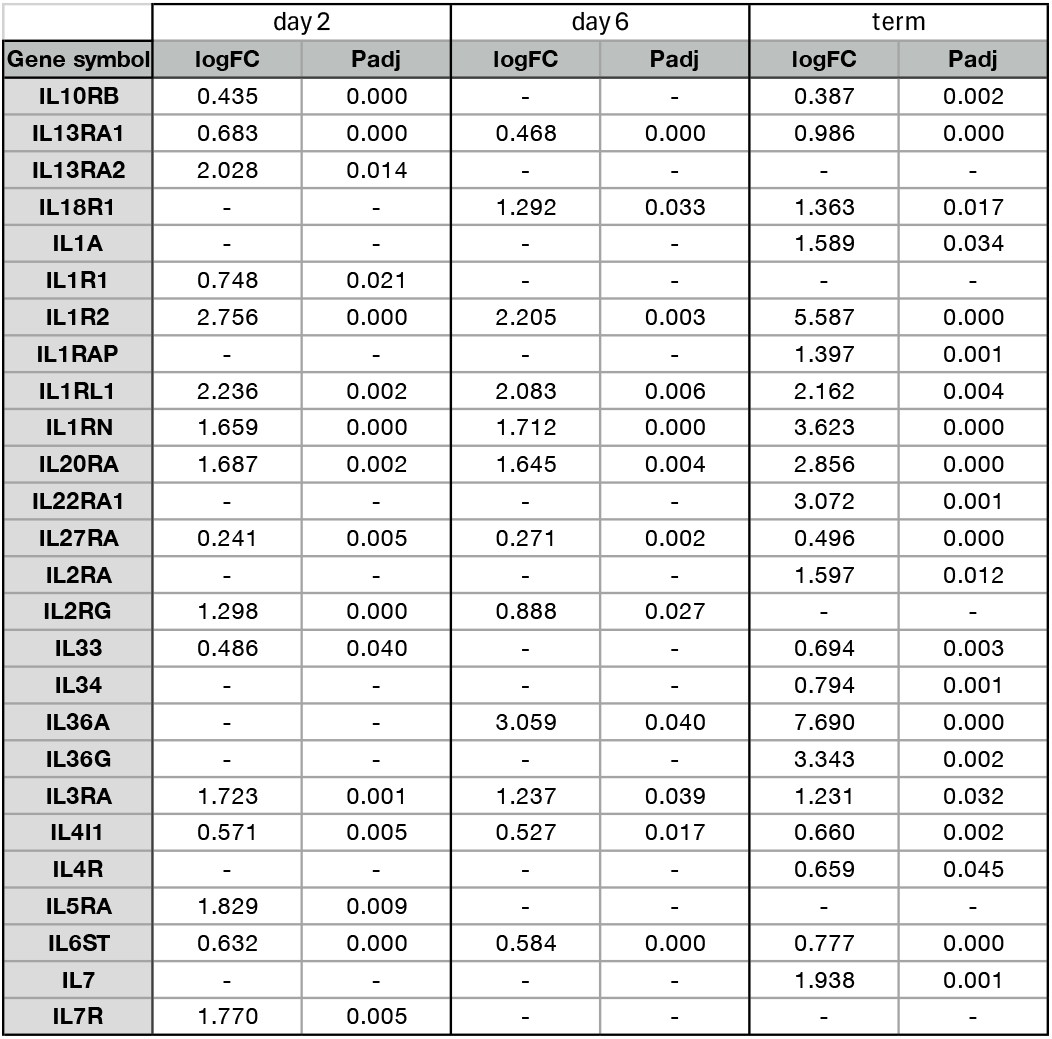
Relative expression of interleukins upregulated in tension conditioned esophageal tissue compared to controls.

### TGFB stimulation of fetal esophageal fibroblasts increases expression of specific collagen subtypes

In order to evaluate the relationship between TGFB signaling and collagen deposition in the esophagus, we stimulated fetal esophageal fibroblasts with TGFB. Following stimulation for varying lengths of time, qRT-PCR was performed to evaluate the expression of COL8A1, COL12A1, and COL6A3. Expression of all three of these collagen subtypes was noted to be significantly upregulated at all timepoints following TGFB stimulation when compared to non-stimulated controls (Fig 8).

**Figure 8.**
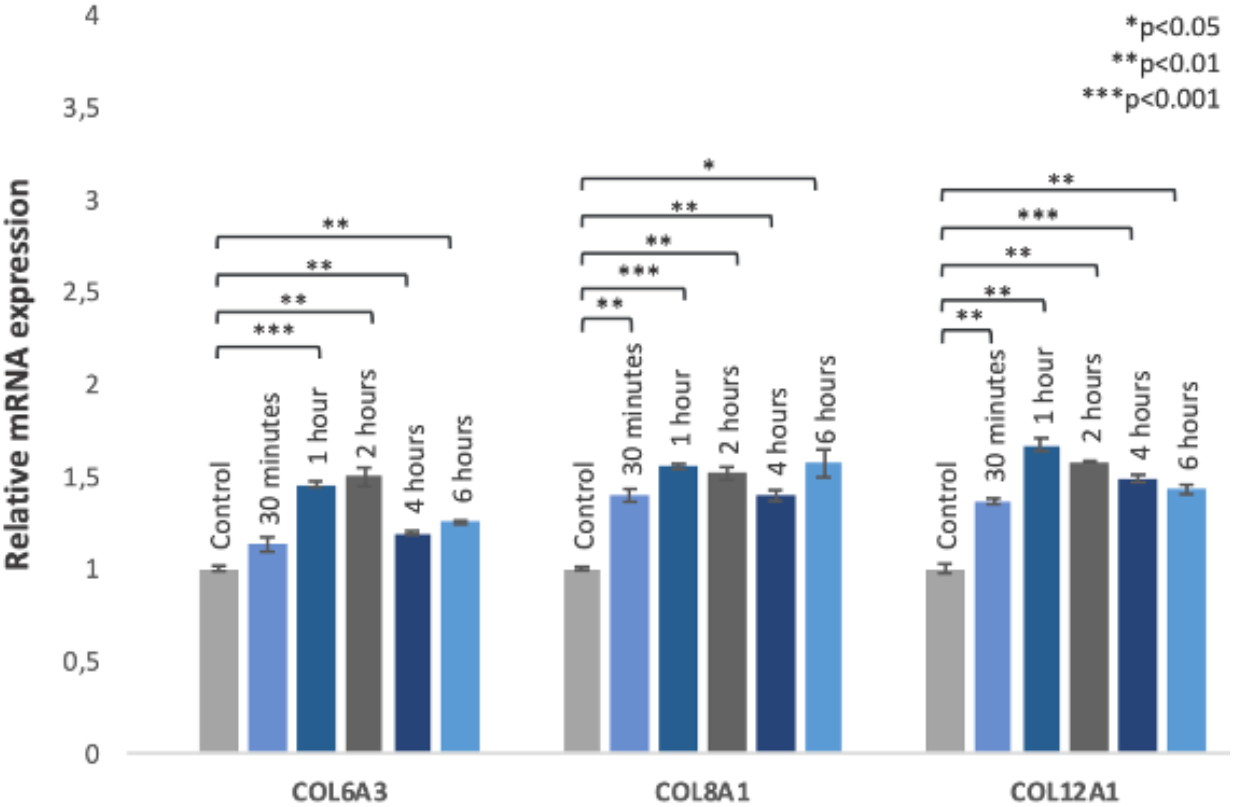
TGFB stimulation of fetal esophageal fibroblast for 30 minutes, 1 hour, 2 hours, 4 hours, and 6 hours. A significant increase in mRNA expression of COL6A3, COL8A1, and COL12A1 is seen at all timepoints compared to control.

## Discussion

In this study, we successfully established a fetal lamb model of LGEA and applied a tension-based lengthening approach to mimic the Foker procedure. Our results demonstrate that esophageal tissue subjected to sustained tension undergoes significant profibrotic changes, as evidenced by upregulation of TGFB signaling, alterations in extracellular matrix organization, and increased collagen deposition. While it is well documented that patients with LGEA have an increased risk of post operative esophageal strictures (8,27), these findings provide the first in vivo proof of the role of tension in conferring a profibrotic phenotype in the tension lengthened esophagus.

The Foker procedure has revolutionized the management of LGEA by utilizing mechanical tension to promote esophageal growth. However, despite its success in achieving primary anastomosis, the prevalence of post operative stricture formation remains (28). Our results suggest that prolonged tension may activate fibroblasts, leading to excessive collagen deposition. The consistent upregulation of specific collagen subtypes (COL8A1 and COL12A1) known to be related to fibrosis (29–32) further supports the hypothesis that tension directly contributes to stricture formation.

Interleukins are known to be important mediators in esophageal fibrosis (33) and future studies will aim to better understand how these inflammatory cytokines are involved in the profibrotic pathway in our model. Further mechanistic studies may elucidate novel targets for anti-fibrotic therapeutics that could be used in conjunction with the Foker procedure to counteract this predisposition for stricture formation. Given that our model recapitulates the fibrosis seen in patients with esophageal stricture, it creates an important opportunity for further translational studies aimed at reversing or preventing these changes.

## Acknowledgements

Research reported in this publication was supported by the National Center for Advancing Translational Sciences of the National Institutes of Health under award number TL1TR001880. The content is solely the responsibility of the authors and does not necessarily represent the official views of the National Institutes of Health.

We gratefully acknowledge the following individuals and organizations for their support and contributions to this work: Aaron Weilerstein, Grace Chau, Marina Heffelfinger, The High Throughput Sequencing Core of the Children’s Hospital of Philadelphia, The CHOP-Penn Proteomics Core, The Pathology Core of the Children’s Hospital of Philadelphia.

## Notes

### Competing Interest Statement

The authors have declared no competing interest.

